# Strategic Leadership Coaching supports Young Executives decision-making

**DOI:** 10.1101/2021.10.10.462360

**Authors:** Charlene Heyns-Nell, Kimberley Clare Williams, David John Hume, Fleur Margaret Howells

## Abstract

Decision-making is central to daily function for executives in any organisation. Strategic leadership coaching (SLC) is an effective way to support complex decision-making, yet empirical neuroscientific data to support is lacking. The purpose of this study was to investigate the effects of SLC on young executive’s cortical arousal and their neural circuitry activation during the completion of computerized tasks which require activation of decision-making circuitry. We hypothesised SLC would improve cortical arousal when engaged with decision-making tasks, specifically increased electroencephalography (EEG) relative alpha band activity and improved neural circuitry engagement, measured as increased amplitude of event-related potential wave components. This study included thirty-one young male executives, of which eighteen underwent 8 sessions of SLC over two months. EEG records were collected thrice from those who underwent SLC (prior, post, and two months post), and twice from the control group (two months apart). The EEG recording session included completion of two decision-making tasks, an Iowa gambling task and Stroop colour-word conflict task. Finding, SLC increased alpha band activity over left frontal and central electrodes, and increased right parietal N170 amplitude and left parietal P300 amplitude. These findings support our hypothesis, as SLC improved cognitive cortical resources (enhanced alpha) which in turn permitted greater efficiency within decision-making circuitry (increased wave component amplitudes). This study provides the first and necessary neurobiological evidence to support and develop this line of research in SLC, and other forms of coaching, as it adds significant value.

## Introduction

Executives are required to engage with complex decision-making daily, they are constantly challenged to restructure and adapt their strategies, to ensure they address the evolution of their respective organization and its market (Eide *et al*., 2020; Uhl-Bien and Arena, 2018). Strategic leadership coaching (SLC) is an effective way to support leaders in complex decision-making processes (Ely *et al*., 2010; Yarborough, 2018). Albeit SLC has been aligned with neuroscience research, the role of SLC in supporting leaders in their decision-making has not yet been validated neuroscientifically.

This study’s aim was to provide neurobiological evidence to the benefits of SLC in young executives. This was achieved by application of non-invasive electroencephalography (EEG) during resting wakefulness and during active task engagement which requires activation of decision-making circuitry. First, we examined changes in arousal, specifically relative alpha frequency activity. Increases in alpha frequency band power have previously been associated with an individual’s readiness to engage with salient stimuli (Hanslmayr *et al*., 2005; Klimesch, 1999; Klimesch *et al*., 2003; Knyazev, 2007). Second, we examined changes in neural circuitry engagement, reflected in event-related potential (ERP) wave component amplitudes, specifically the N170 and P300. Where enhanced wave component amplitudes, reflect greater recruitment and efficiency of engagement in decision-making processes (Kok, 2001; Saliasi *et al*., 2013). First we postulated, relative alpha frequency band power will increase with SLC, as coaching will improve cortical resource availability. Second, with improved cortical resources the recruitment and efficiency of specific neural circuitry engagement, wave-component amplitudes, involved in decision-making processes would be enhanced.

## Materials and Methods

### Research design

This study included two groups of participants, an intervention group (IG) and control group (CG). The IG’s electroencephalography (EEG) was recorded at three time points, over 4 months. Time point 1, was prior to initiating the 8 sessions of SLC. Time point 2, was within one week post 8 sessions of SLC, being a 2 month period. Time point 3, was 2 months post SLC. The CG’s EEG was recorded at two time points, with two months between testing to mimic the IG group’s time point 1 and 2, **Figure 1**.

**Figure 1.**
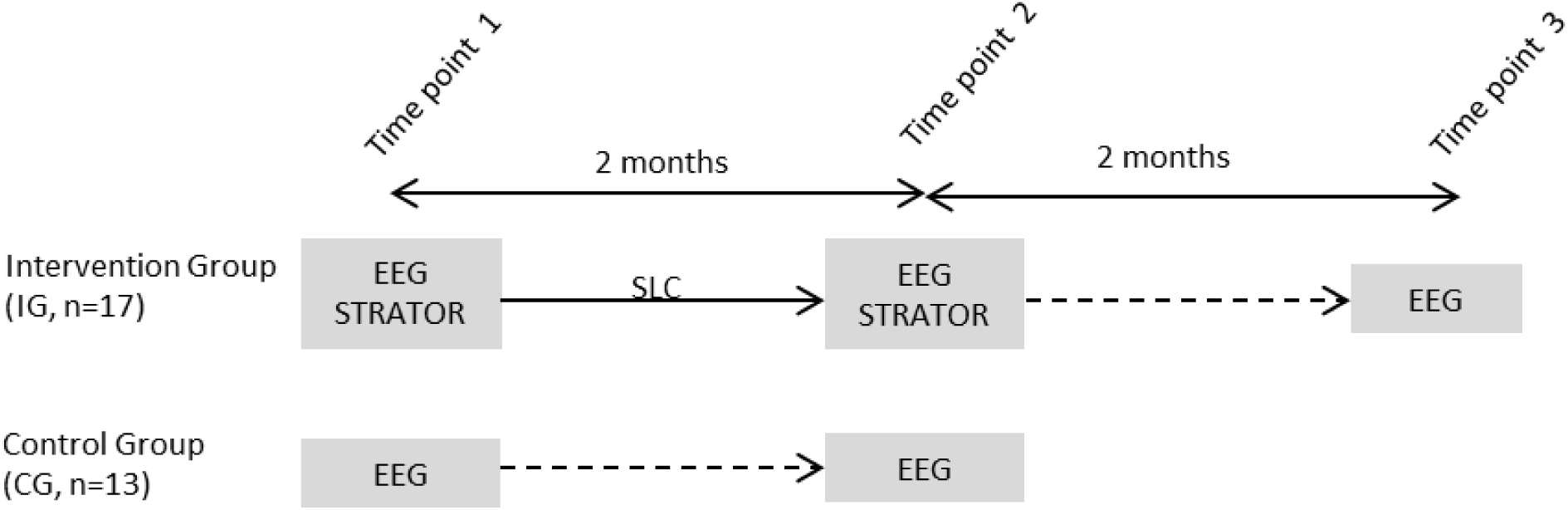
This study included two groups of participants, an Intervention Group (IG) and Control Group (CG). The IG included 17 young executives; they completed 8 sessions of Strategic Leadership Coaching (SLC) over a 2 month period. The IG group completed three electroencephalography (EEG) recordings, prior to the initiation of coaching (time point 1), immediately post coaching (time point 2), and two months post coaching (time point 3). The IG group completed the STRATOR scale prior and post strategic leadership coaching. The CG included 13 young executives who completed two EEG recordings (time point 1 and time point 2), these records were 2 months apart.

### Participants

We recruited young executives between the ages of 25-45 via social media and in-house company secretaries, using established networks held by our team. For this study, we recruited a convenience sample, which led us to excluding suitable female participants from this primary study. Inclusion to the study required each participant to be top, senior or middle managers, who held accountability for strategic decision-making in their organisation, with a comparative salary bracket related to this level of responsibility. Participants were also required to hold general knowledge and overall awareness of their organisation’s policies and day-to-day functioning, and be able to access their organisation’s financial and strategic information. Exclusion criteria included that which may have affected EEG record: any form of epilepsy/seizure disorder in self or family history thereof; significant brain trauma or surgical intervention.

### Strategic Leadership Coaching (SLC)

The IG participants received eight one-hour sessions of SLC over a two-month period. The first objective was to create a coaching presence, through building rapport and confidentiality, achieved by completing a ‘coaching contract’. The second objective was to establish the strategic orientation of the participant within their organization, achieved by administering the STRATOR scale, see section 2.4.1, and asking powerful open-ended questions linked to this scale. Throughout the coaching sessions reference to neuropsychological models which are involved in the development of strategy, objective setting and complex decision making were woven in. One such model employed was the diffusion model which supports effective decision-making. To support participants in creating their vision, strategic planning, and supporting process flows the funneling technique was applied. As with the STRATOR, powerful open-ended questions and active listening, while the coach maintained an unbiased view, this is essential to the success of participant’s coaching engagement. Several coaching models were applied, ARAA and OSCAR, to facilitate participants to engage with their company’s vision, and how, by implementing various strategic initiatives, they could introduce greater process flow and unity within their teams. In addition, throughout all coaching sessions, the International Coach Federation (ICF) core competencies of setting the foundation, co-creating relationships, communicating effectively and cultivating learning and growth were incorporated.

### STRATOR scale

The IG completed the STRATOR prior- and post-the 8 coaching sessions. The STRATOR scale (Heyns-Nell, 2009) was developed to assess individual strategic orientation in their respective organisation. This 53-item scale assesses management emphasis on: Strategic Orientation (A, 46 items), Business Performance in relation to strategic orientation (B, 3 items), and Top Management Emphasis on organisation’s strategic orientation (C, 4 items). The STRATOR employed a 6-point Likert scale (5 = strongly agree, 4 = agree, 3 = neutral, 2 = disagree, 1= strongly disagree, 0 = don’t know). Strategic Orientation (A) included five scales, comprising 46 items. (A1) Market Orientation, the degree to which companies obtain customer information and use it for strategic planning, comprised of 20 items, divided into three sub-scales: (A1a) Strategic implementation with 8 items, measures the degree to which a business unit obtains and uses information from customers; (A1b) Strategic development with 8 items, measures the development of strategies to meet customer needs; (A1c) Use of customer information with 4 items, measures the response to customer needs and wants within their market. (A2) Corporate Entrepreneurship comprised of 6 items, measures the understanding of a company’s appetite for business venturing within western philosophy, including pro-activeness, risk taking, and innovativeness. (A3) Innovativeness, comprised of 4 items, measures the design and execution of new ideas, products and processes, e.g. being first to market with new products and/or refinement of existing products. (A4) Learning Orientation comprised of 12 items, which measures the overall organisational culture in terms of its commitment to learning, with three subscales: (A4a) Commitment to Learning with 4 items, a measures the shared value in understanding the causes and effect of their actions; (A4b) Open-Mindedness with 4 items, measures the ability of employees to challenge the status quo and seek improved ways of doing things; (A4c) Shared Vision with 4 items, measures the ownership that employees take in regards to the goals and values that are reflected in the company’s vision. And (A5) Technology Orientation comprised of 4 items, which measures a company’s ability to acquire and use research to develop technical solutions for customers. Then the remaining two sub-scales, Business Performance (B) with 3 items measured impact of strategic orientation, including sales, market share, and profitability, and Top Management Emphasis (C) with 4 items measured the support by top management through their engagement with market trends and competitive activities.

### Levenson’s Self-Report Psychopathy Scale (LSRP)

The LSRP (Levenson *et al*., 1995) was developed to assess primary and secondary psychopathy. This scale was included, to manage the potential influence of psychopathy on EEG frequency band activity. Psychopathy is suggested to present frequently in business leaders (Boddy, 2011) and has been associated with decreased alpha band power (Calzada-Reyes *et al*., 2013; Ortega-Noriega *et al*., 2015; Tillem *et al*., 2019), the frequency of interest in this study. The LSRP consists of 26 items, answered on a 4 point Likert scale from 1 (*strongly disagree*) to 4 (*strongly agree*), designed to assess similar domains as the Psychopathology Checklist (Schroeder *et al*., 1983). The first domain refers to a callous, manipulative, and selfish use of others (e.g., “Success is based on survival of the fittest”; “I am not concerned about the losers” and “For me, what’s right is whatever I can get away with”). The 2^nd^ domain is concerned with impulsivity and poor behavioural controls (e.g., “I find myself in the same kinds of trouble, time after time” and “I am often bored”) (Lynam *et al*., 1999; Miller *et al*., 2008).

### Iowa gambling task (IGT)

The IGT was designed to assess risk preferences by stimulating real-life decision making by incorporating uncertainty, rewards, and penalties (Bechara *et al*., 1997). The version employed in this study presented the participant with four different tokens (diamond, circle, square, crystal), versus the traditional card decks, presented in the same order in a square formation on a computer screen. Players are instructed to select a token, the participant is then presented with their accumulated value after the selection of a token, where the traditional version provided a monetary value. The participant was informed that each token had both rewards and penalties associated with it, prior to starting the task. Two indices were calculated from the IGT. The *long-term consequences index*, reflected the selection of advantageous tokens, being the selection of either the circle or square, as they yielded a positive outcome overall. Where total value for circle amounted to 15,700 and square amounted to 12,570, while selection of crystal amounted to −50940 and diamond amounted to −22650. Then *index for bias to infrequent loss*, were selection of circle and crystal tokens, respectively their loss occurred 8.6% and 22.6%, while the square was 38% and diamond 47.3%. Values for these two indexes and *total scores* were extracted per 100 trials and overall, being the outcome of 300 trails.

### Stroop colour-word conflict task

The Stroop colour-word conflict task produces cognitive interference when a decision needs to be made and response is made. Our study’s task was an adaptation from the original Stroop task (Stroop, 1935). The version employed in this study included the presentation of 60 colour-word conflicts, using four colours (yellow, green, red, blue; 15 of each colour), the participant was required to respond to the colour of the word, not the written word, as quickly and as accurately as possible. To maintain the colour-word conflict, i.e. Stroop effect, the participant was presented intermittently with 20 greyed words (yellow, green, red, blue; 5 of each colour), in these instances the participant was to respond to the written word and not the colour grey. In our version of the task we also activated working memory, by inserting 20 grey squares and participants were asked to count the number of squares presented during the task. Accuracy, omissions in response, and response times were extracted for the 60 colour-word conflicts.

### Electroencephalography (EEG)

EEG recordings were taken during resting states and during the completion of the two decision-making tasks. The resting states included 3 minutes of resting eyes open, where the participant maintained focus on a central fixation cross on the computer screen, and 3 minutes of resting eyes closed, maintaining the same position, sitting relaxed in front of a computer screen. For all EEG records digital tagging was sent from Eprime version 2.0 (Psychology Software Tools Inc, 2012) to software collecting the EEG record, Acqknowledge version 4.1, this ensured discrete windows for each record and stimulus-time-locked extraction of ERPs.

A simple EEG montage that included bilateral frontal (F_3_ and F_4_), central (C_3_ and C_4_) and parietal (P_3_ and P_4_) electrodes was used. With standard 10/20 montage EEG caps (Electro-Cap International, Inc.), of either medium or large size depending on head circumference of participant. Participants were grounded peripherally, linked earlobe reference was applied, and electrooculography (EOG) was recorded. The EEG system used was the Biopac MP150 system with 100C EEG amplifiers and EOG amplifier (Biopac Systems, Inc.). Digital EEG data and analogue data, from E-prime, were collected via the MP150 system, with a sampling rate of 500 Hz, and were visualised real-time using Acqknowledge 4.1 (Biopac Systems, Inc.). For EEG data processing, data were eye blink and movement corrected, using automated ICA correction in Acqknowledge 4.1, band pass filtered 0.1–30 Hz, then Fourier transformed, using an in-house Matlab GUI. Relative alpha frequency band power was extracted for resting states and cognitive tasks, calculated by extracting absolute alpha frequency band power (7-14 Hz) and dividing it by the sum of absolute power extracted (0.1-30 Hz), multiplied by 100.

Event-related potentials (ERPs) were extracted for colour-word conflict stimuli from the Stroop task. Individual ERPs were extracted (−200-500msec) with automated artefact rejection of +100 μvolts and –100 μvolts, a grand mean average ERP for each participant was determined and baseline corrected (−200-0msec), and visually inspected. Robust wave components were identified and windows applied to extract latency to peak amplitude and peak amplitude. Three wave components were extracted: P150 for frontal electrodes with a window of 120-220msec, N170 for central and parietal electrodes with a window of 120-220msec, and P300 for frontal, with a window of 220-450msec, then central and parietal electrodes, with a window of 220-400msec.

### Statistical analysis

All analyses were performed in Statistica version 13.5 (TIBCO Software Inc., 2016). All data were subjected to distribution checking, with application of Shapiro-Wilks W test, which guided the analysis steps, i.e. data of normal distribution underwent parametric statistics and data not of normal distribution underwent non-parametric statistics.

### Group differences

Analysis of the two groups (IG and CG) over time (time point 1 and time point 2) the following statistical approach was applied. For data where distribution was not normal, to determine group differences Mann-Whitney-U test was applied, to determine differences over time Wilcoxan matched pairs test by group was applied. For data that were of normal distribution repeated measures ANOVA were applied which provided the interaction of time and groups comparisons in one model, followed by Bonferroni correction post-hoc analyses. Two group correlation analyses were applied in accordance with distribution of data, Pearson’s for parametric and Spearman’s for non-parametric. Correlations were performed within a task, investigating behavioural performance with EEG parameters, then behavioural and EEG parameters with LSRP scale. *A priori*, to account for the number of correlations performed, threshold for reporting significant correlations was set greater than ±0.70, with p-value < 0.001.

### Within the intervention group (IG)

STRATOR scale data was collected at time point 1 and time point 2 for IG. The data were investigated for differences over time, as appropriate. For data where distribution was normal dependent T-tests were performed, and when data was not of normal distribution Wilcoxan matched pairs test was performed. Analysis of IG over three time points, prior (time point 1), post (time point 2), and two months follow-up (time point 3) the following statistical approach was applied. For data where distribution was not normal, to determine differences between time points Chi Square tests were performed, when significance was apparent it was followed with Wilcoxan matched pairs test. For data that were of normal distribution dependent T-tests were performed. Within IG correlation analyses were applied in accordance with distribution of data, Pearson’s for parametric and Spearman’s for non-parametric. Correlations were performed between the Strator scale and behavioural and EEG parameters, for time point 1 and time point 2. A priori, to account for the number of correlations performed, threshold for reporting significant correlations was set greater than ±0.70, with p-value < 0.001.

## Results

### Population characteristics

Participants’ age and BMI did not differ between the intervention group (IG_(n=18)_ age=35.44±4.03, BMI=27.55±4.94) and control group (CG_(n=13)_ age=35.33±6.00, BMI=27.48±5.20). There were two left handed participants in IG and one in the CG. Two participants in each of the groups reported being smokers. Chronic medication was prescribed to five participants in the IG, as follows: (1) cholesterol lowering (Aspavor 30mg/d; active ingredient atorvastatin a HMG-CoA reductase inhibitor) and gout medication (Pericos 300mg/d; active ingredient allopurinol an inhibitor of the enzyme xanthine oxidase); (2) cholesterol lowering medication (Crestor 10mg/d, active ingredient rosuvastatin another HMG-CoA reductase inhibitor); (3) acid reflux medication (Nexium 20mg/d; active ingredient esomeprazole a proton pump inhibitor); (4) antihistamine (Loratadine; an inverse agonist of peripheral histamine H_1_ receptors); (5) steroid nasal spray (Nexomist, active ingredient mometasone furoate which reduces inflammation) used as needed by the participant. Chronic medication was prescribed to three participants in the CG, as follows: (1) insulin to manage type II diabetes as necessary; (2) antidepressant (escitalopram 10mg/d, a selective serotonin reuptake inhibitor); (3) acute 3 month antidepressant medication regime, at his first visit he was taking an anxiolytic for 7 days (Diazepam 5mg/d) with an antidepressant for three months, ergo at his second visit he was taking antidepressant medication (Thaden 7.5mg/d, active ingredient dosulepin a tricyclic antidepressant).

### Group differences

#### Resting state activity

For resting state activity only during resting eyes open there was an increase in relative alpha over time for right frontal electrode, this was evident for the population (F_4_ z_(n31)_=2.52, p=0.011) and then for IG (F_4_ z_(n18)_=2.02, p=0.042), **Table 1**.

**Table 1.**
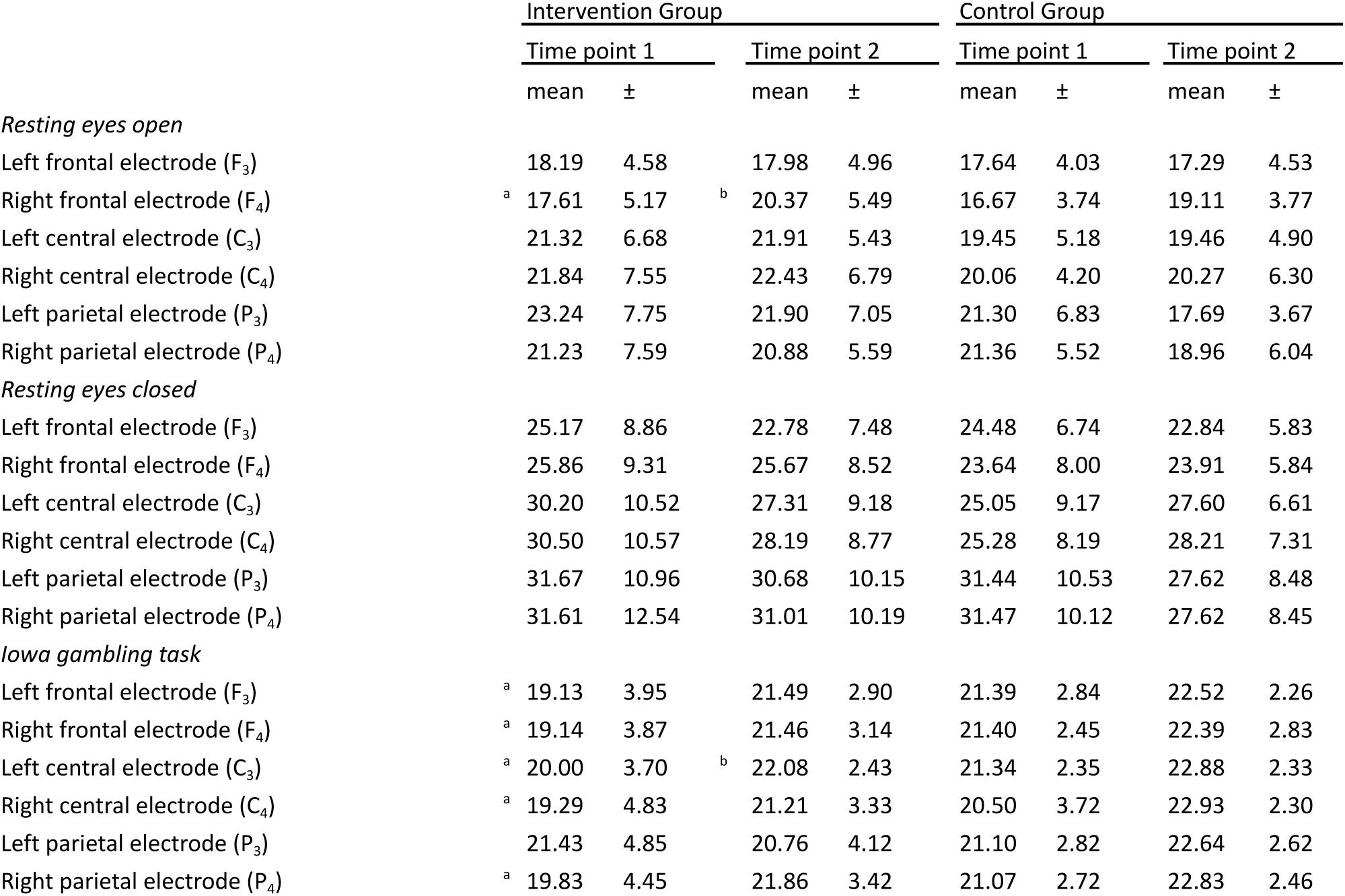
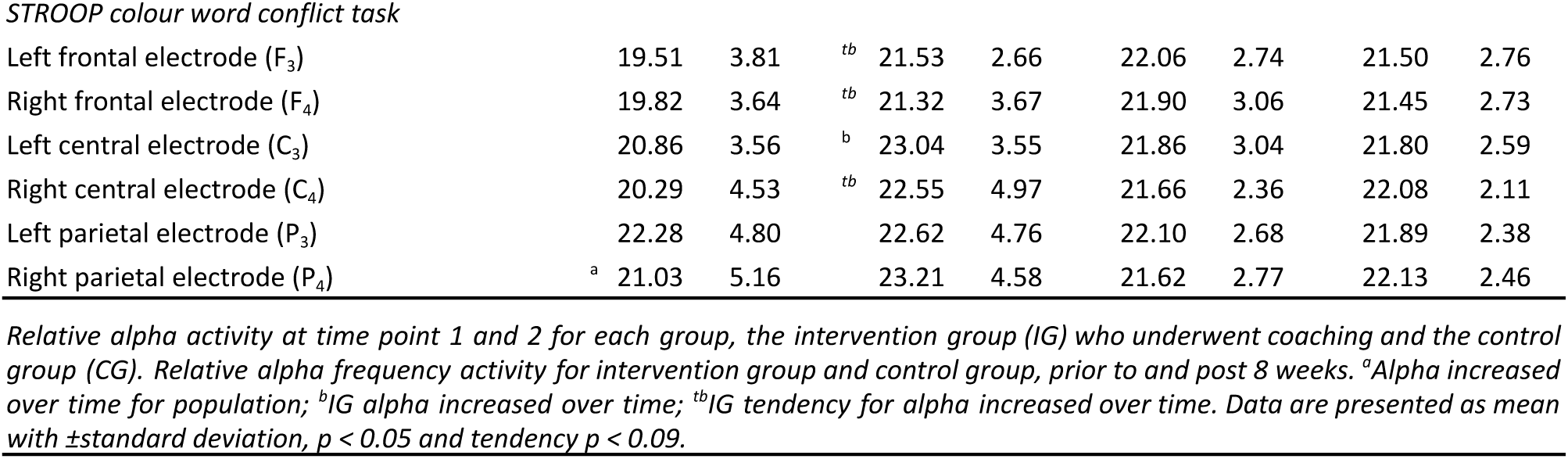
Relative alpha activity of the two groups over 2 time points

#### Iowa gambling task

Behaviourally, IGT performance yielded a group difference at the first time point, where long-term consequence index for the first 100 trails was higher for CG when compared to IG (z_IG17,CG12_=-2.05, p=0.039). Overtime our population’s IGT long term consequences improved over the 300 trials (z_n28_=3.30, p=0.00096) and for each bin of 100 trials (1^st^ 100 trials z_n28_=3.18, p=0.0014; 2^nd^ 100 trials z_n28_=3.46, p=0.00053; 3^rd^ 100 trials z_n28_=2.65, p=0.0080). Then bias to infrequent loss was, during the 1^st^ 100 trials, also found to improve over time for our population (z_n27_=2.06, p=0.038). When investigating the individual groups, improvement for IGT long term consequences were limited to IG, over the 300 trails (z_n17_=3.14, p=0.0016) and for each bin of 100 trials (1^st^ 100 trials z_n17_=2.72, p=0.0064; 2^nd^ 100 trials z_n17_=3.10, p=0.0019; 3^rd^ 100 trials z_n17_=2.61, p=0.0089), **Table 2**.

**Table 2.** Behavioural performance of the two groups over 2 time points

|  | Intervention Group |  |  |  | Control Group |  |  |  |
| --- | --- | --- | --- | --- | --- | --- | --- | --- |
|  | Time point 1 |  | Time point 2 |  | Time point 1 |  | Time point 2 |  |
| | mean | ± | mean | ± | mean | ± | mean | ± |
| *Iowa gambling task* |  |  |  |  |  |  |  |  |
| *Long-term consequence* |  |  |  |  |  |  |  |  |
| First 100 trials | <sup>a</sup> <sup>b</sup> 0.90 | 0.26 | <sup>c</sup> 4.20 | 11.36 | 1.10 | 0.30 | 1.90 | 1.82 |
| Second 100 trials | <sup>b</sup> 1.20 | 0.50 | <sup>c</sup> 7.70 | 15.53 | 1.60 | 1.46 | 8.30 | 15.87 |
| Third 100 trials | <sup>b</sup> 6.10 | 12.87 | <sup>c</sup> 8.30 | 16.08 | 2.10 | 2.32 | 9.90 | 18.93 |
| All 300 trials | <sup>b</sup> 1.20 | 0.52 | <sup>c</sup> 5.10 | 11.62 | 1.40 | 0.80 | 2.90 | 3.51 |
| *Bias to infrequency loss* |  |  |  |  |  |  |  |  |
| First 100 trials | <sup>b</sup> 1.60 | 0.57 | 4.70 | 11.12 | 1.70 | 0.81 | 2.00 | 1.19 |
| Second 100 trials | 1.90 | 1.25 | 4.50 | 7.28 | 2.20 | 0.99 | 2.40 | 1.91 |
| Third 100 trials | 1.90 | 4.40 | 3.00 | 5.31 | 5.90 | 13.61 | 6.60 | 14.10 |
| All 300 trials | 3.00 | 0.92 | 3.70 | 7.22 | 2.00 | 0.92 | 2.40 | 1.87 |
| *Score* |  |  |  |  |  |  |  |  |
| First 100 trials | -5779 | 5745 | -5269 | 6644 | -6971 | 3209 | -4461 | 3863 |
| Second 100 trials | -2272 | 4128 | -2491 | 6451 | -4295 | 3761 | -254 | 4089 |
| Third 100 trials | -4758 | 5885 | -3146 | 6272 | -3418 | 7071 | -2496 | 6239 |
| All 300 trials | -12810 | 10586 | -10908 | 15155 | -14685 | 7314 | -7212 | 12213 |
| *Stroop colour-word conflict* |  |  |  |  |  |  |  |  |
| Response time (ms) | 848 | 230 | <sup>c</sup> 823 | 187 | 986 | 219 | 999 | 237 |
| Omissions (%) | 0.28 | 0.86 | 0.19 | 0.54 | 0.51 | 1.25 | 1.92 | 5.96 |
| Inaccurate responses (%) | 1.57 | 2.25 | 1.76 | 2.39 | 0.64 | 0.84 | 1.41 | 2.53 |
| Square count | 19.38 | 1.19 | 19.77 | 0.64 | 19.69 | 2.35 | 19.07 | 2.84 |

*Behavioural performance outcomes at time point 1 and 2 for each group, the intervention group (IG) who underwent coaching and the control group (CG). <sup>a</sup>Time point 1 CG > IG. <sup>b</sup>Time point 1 to time point 2 improvement was found for full population. <sup>c</sup>IG improved from time point 1 to time point 2. Data are presented as mean with ± standard deviation, $p < 0.05$ .*

Relative alpha activity, during the IGT increased overtime for the population for bilateral frontal electrodes (F_3_F_(1,29)_=6.908, p=0.013; F_4_F_(1,29)_=4.602, p=0.027), bilateral central electrodes (C_3_Z_(n31)_=2.70, p=0.0068; C_4_F_(1,29)_=7.74, p=0.0093), and right parietal electrode (P_4_F_(1,29)_=8.751, p=0.0061). Then IG showed increased relative alpha activity for left central electrode over time (C_3_z_(n18)_=2.19, p=0.027), **Table 1**.

#### Stroop colour-word conflict task

Behaviourally, the IG group overtime showed shorter response times for the Stroop colour-word conflict task (F_(1,28)_=5.12, p=0.031), **Table 2**. Relative alpha activity, during the Stroop colour-word conflict task, increased overtime for the population for right parietal electrode (F_(1,29)_=5.22, p=0.029, TI to T2 p=0.016). Then IG showed increased relative alpha activity for left central electrode over time (C_3_z_(n18)_=2.19, p=0.027), with a similar yet weak tendency for increased relative alpha activity for right central electrode (C_4_z_(n18)_=1.67, p=0.093), then tendencies for increased relative alpha activity for bilateral frontal (F_3_F(1,29)=4.524, p=0.042, post-hoc IG p=0.093; (F_4_z_(n18)_=1.76, p=0.077)), **Table 1**. ERP wave components, from time point 1 to time point 2, IG showed increased P300 amplitude for the left parietal electrode (P_3_z _n18_=2.23, p=0.019), and a tendency towards increased N170 amplitude over the right parietal electrode (P_4_z_n18_=1.93, p=0.052). While CG showed decreased amplitude over the right central electrode for event-related potential wave form P300 (C_4_z _n12_=2.19. p=0.028), **Figure 2**.

**Figure 2.**
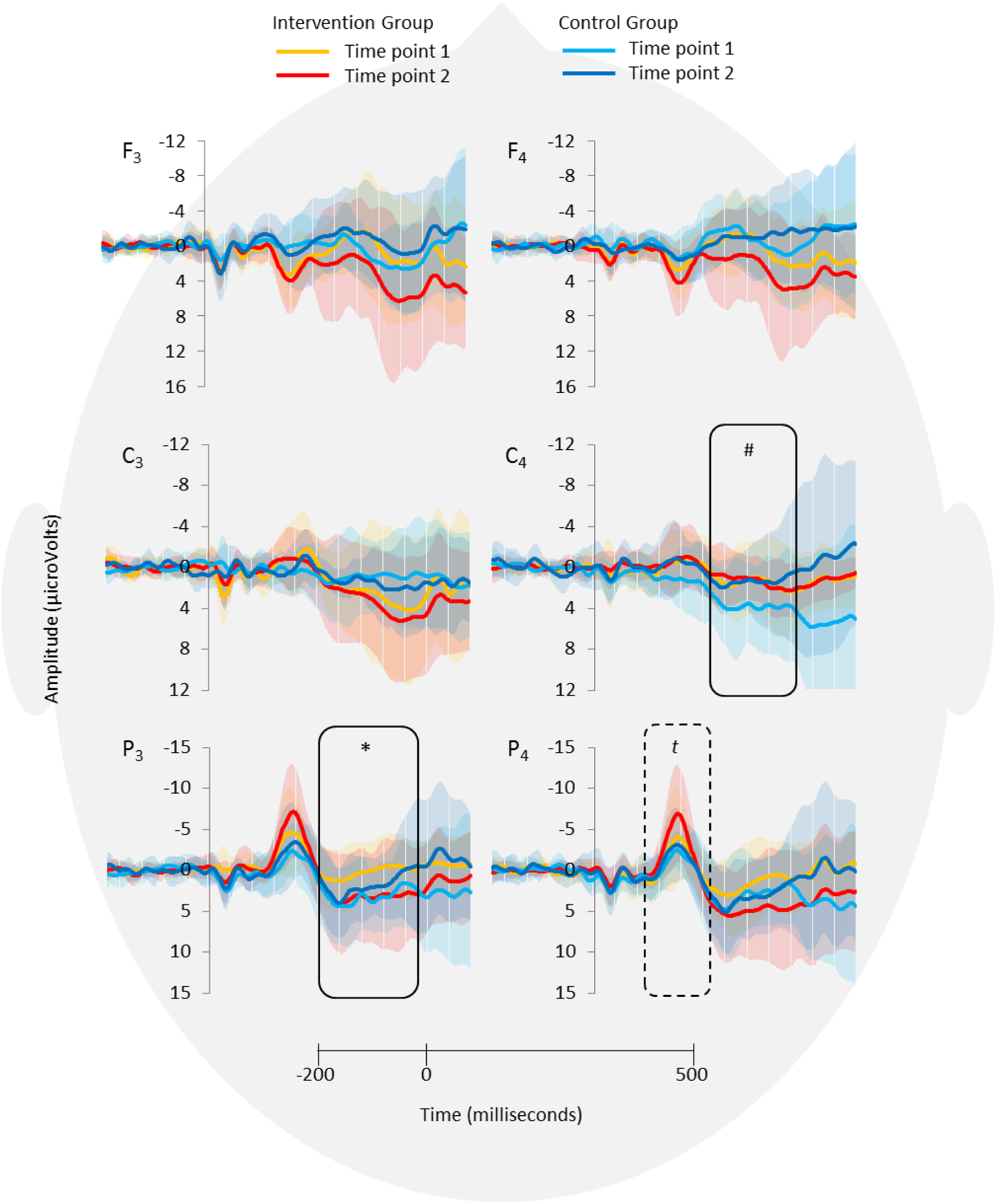
Stroop colour-word conflict event-related potentials (−200 to 500msec) for the Intervention Group (IG_n18_) and Control Group (CG_n12_), at time point 1 and time point 2, being two months apart and where IG group has undergone 8 sessions of Strategic Leadership Coaching (SLC). Event-related potentials are shown for left and right frontal (F_3_, F_4_), central (C_3_, C_4_), and parietal (P_3_, P_4_) electrodes. *IG showed increased P_3_ P300 amplitude over time, and a ^*t*^tendency towards increase for P_4_ N170 amplitude, while ^#^CG showed decreased C_4_ P300 amplitude. Data are presented as grand mean averages for each group with error bars reflecting their standard deviation.

#### Levenson Self-Report Psychopathy scale (LSRP)

Scale revealed no group differences in either domain, where the 1^st^ domain refers to callous, manipulative, and selfish use of others, and 2^nd^ domain refers to impulsivity and poor behavioural control (Lynam *et al*., 1999; Miller *et al*., 2008). Our participants when compared to normative data were found to present with lower self-report psychopathy symptomology (IG_(n18)_ 1^st^ domain 25.13±6.45, 2^nd^ domain 15.90±2.61; CG_(n13)_ 1^st^ domain 23.13±4.04, 2^nd^ domain 16.00±4.09), where normative data on males, report 1^st^ domain score of 32.96 and 2^nd^ domain scored of 20.04 (Levenson *et al*., 1995).

The primary domain of the LSRP yielded significant relationships with relative alpha frequency activity at time point 1. IG’s primary domain of the LSRP correlated negatively with relative alpha, during resting eyes closed for bilateral central electrodes (REC C_3_ R_Spearman’s(n18)_=-0.72, p=0.00066; REC C_4_ R_Spearman’s(n18)_=-0.72, p=0.00072), **Figure 3**.

**Figure 3.**
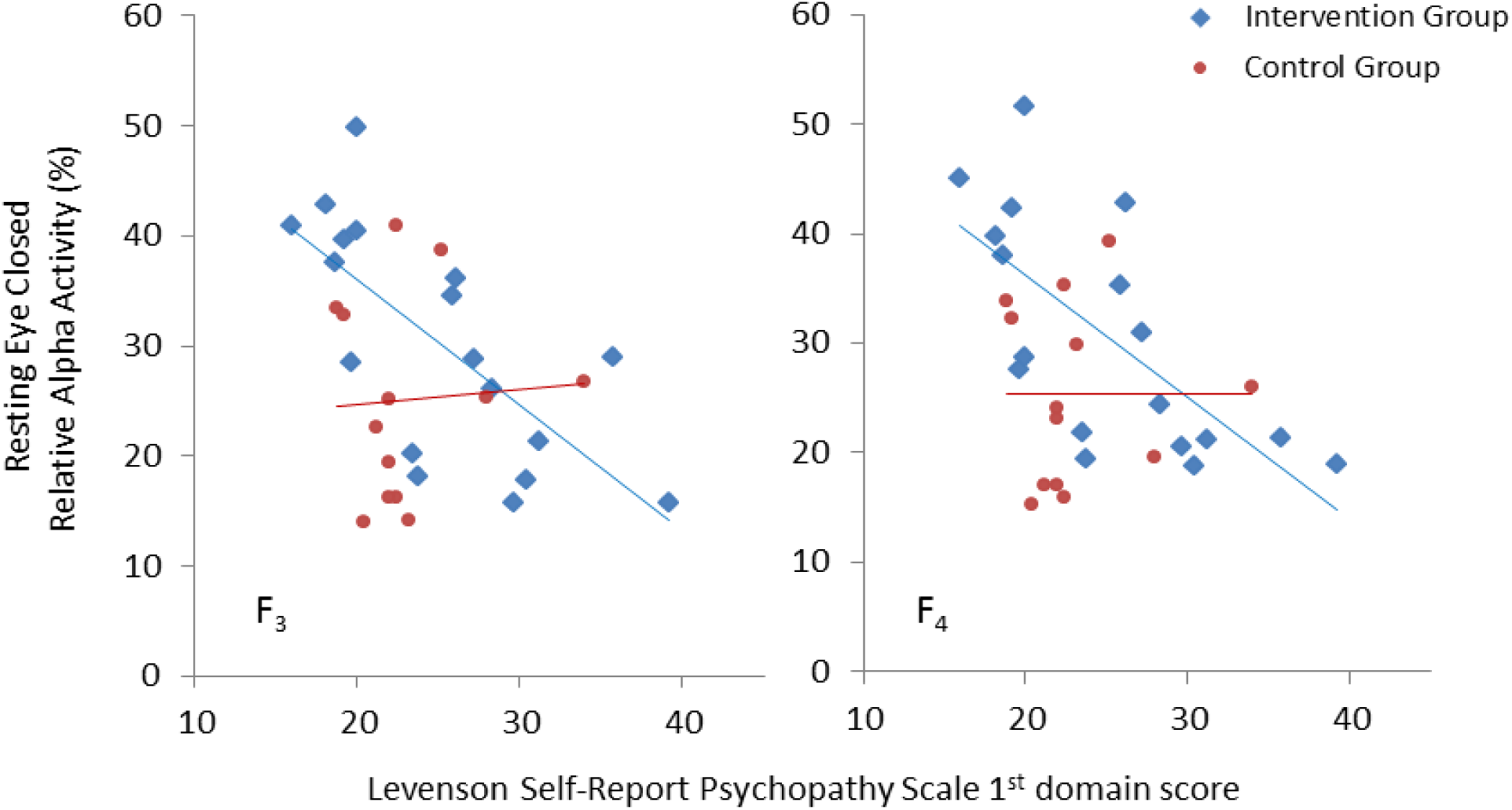
Relative alpha frequency activity (%) during resting eyes closed at time point 1 negatively correlated with Levenson Self-Report Psychopathy Scale’s 1^st^ domain score for Intervention Group, This was apparent for left frontal electrode (F_3_) and right frontal electrode (F_4_). Individual data points are shown for each participant with linear trend line to reflect slope of correlation.

### Within the Intervention Group

#### STRATOR

The IG_(n=18)_ were administered the STRATOR prior and post the intervention. Then Market Orientation’s sub-category, Use of Customer Information: “Payment terms or credit policies with clients take their needs into account” decreased over time (z=1.98, p=0.046). Aspects of Learning Orientation increased over time, under subcategory Commitment to Learning: “Management sees survival and competitiveness dependent on the ability to learn” (z=2.10, p=0.0.35) and under subcategory Open-Mindedness: “Not afraid to question the shared assumptions made about approach to business” (z=1.96, p=0.049). And under Top Management Emphasis “We must gear up to meet customers’ future needs” decreased over time (z=2.02, p=0.043), **Table 3**.

**Table 3.**
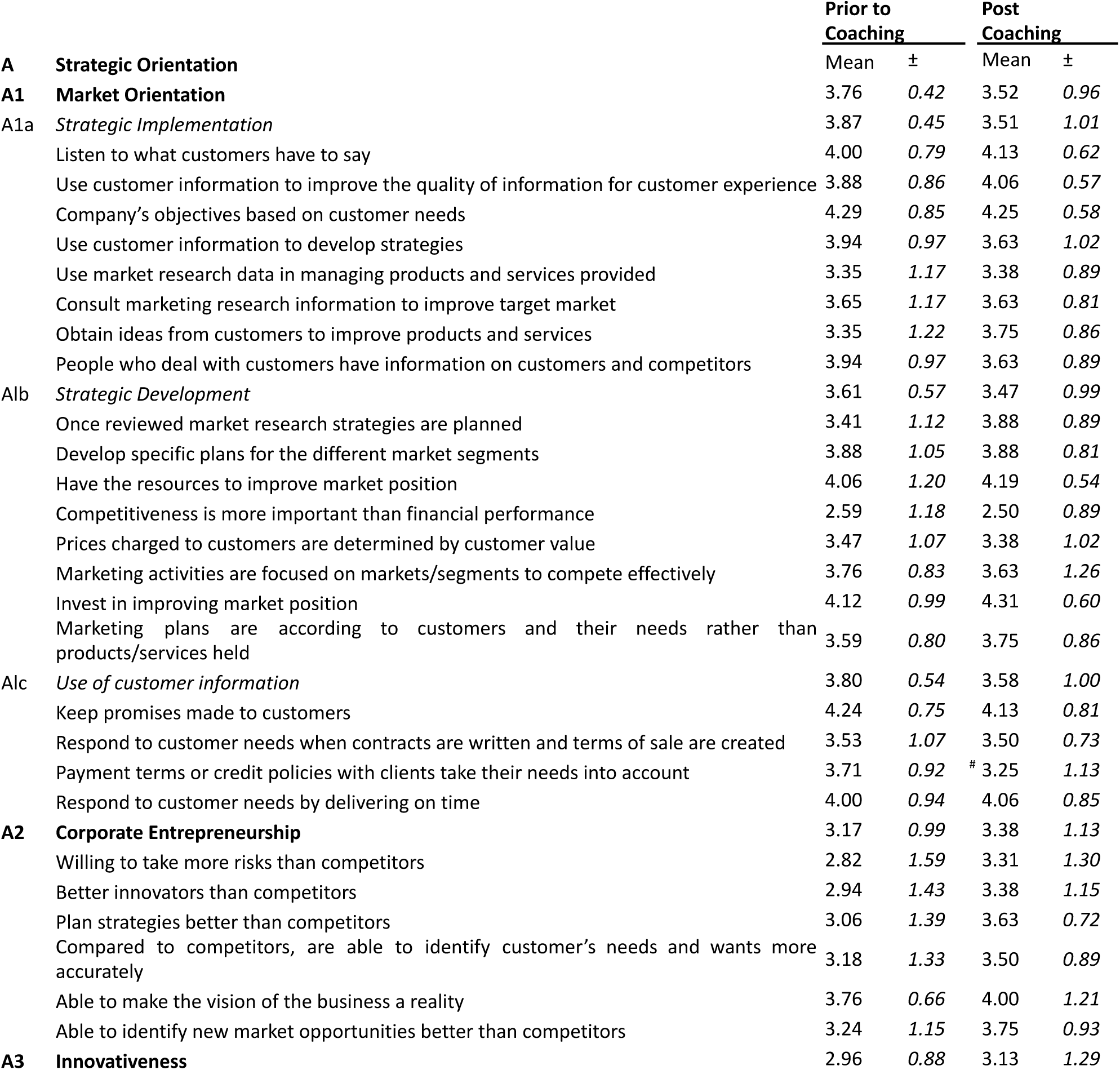
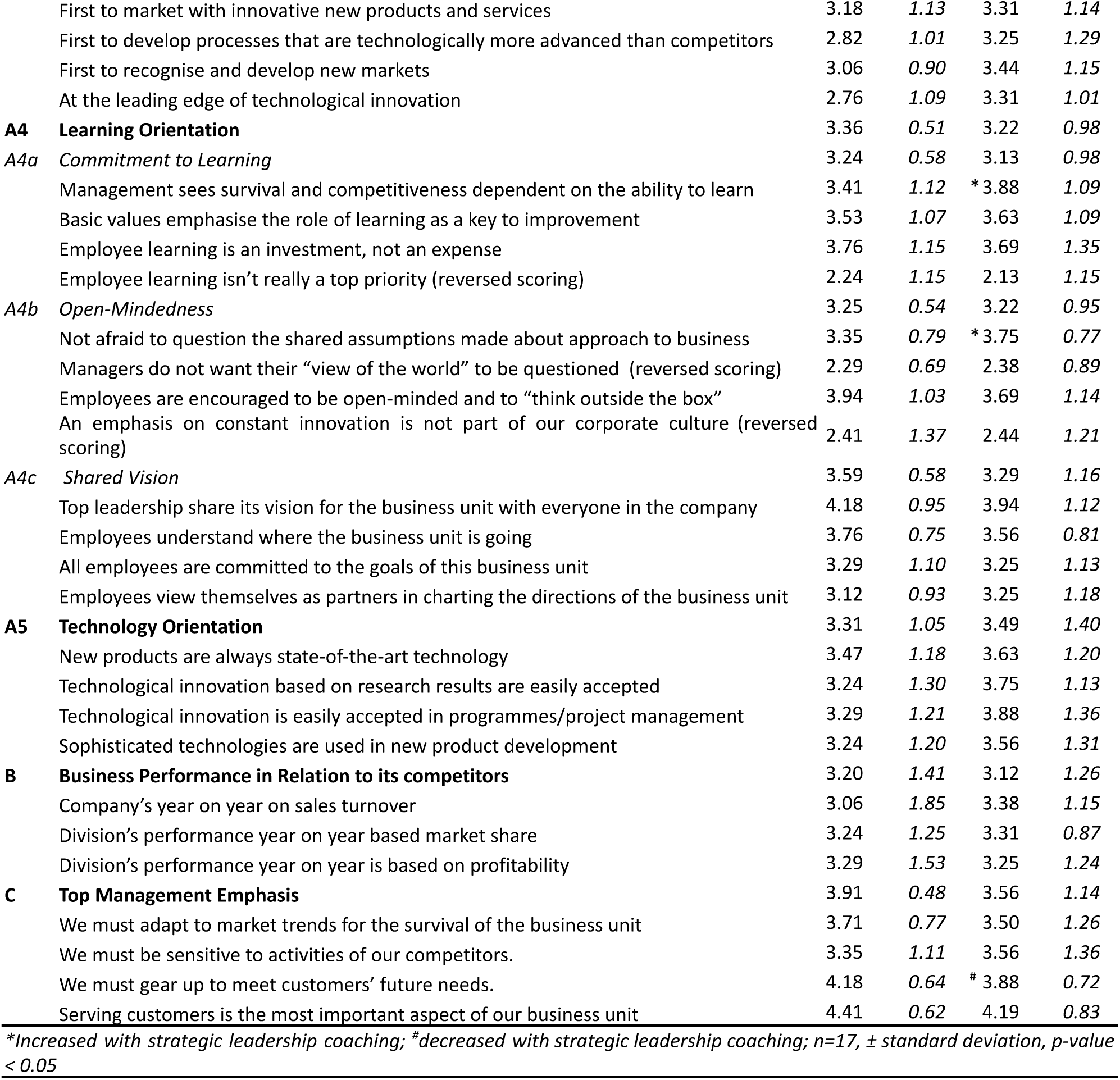
STRATOR for the Intervention Group over 2 time points

|  | **Prior to Coaching** |  | **Post Coaching** |  |
| --- | --- | --- | --- | --- |
|  | Mean | ± | Mean | ± |
| **A Strategic Orientation** |  |  |  |  |
| **A1 Market Orientation** | 3.76 | 0.42 | 3.52 | 0.96 |
| A1a *Strategic Implementation* | 3.87 | 0.45 | 3.51 | 1.01 |
| Listen to what customers have to say | 4.00 | 0.79 | 4.13 | 0.62 |
| Use customer information to improve the quality of information for customer experience | 3.88 | 0.86 | 4.06 | 0.57 |
| Company's objectives based on customer needs | 4.29 | 0.85 | 4.25 | 0.58 |
| Use customer information to develop strategies | 3.94 | 0.97 | 3.63 | 1.02 |
| Use market research data in managing products and services provided | 3.35 | 1.17 | 3.38 | 0.89 |
| Consult marketing research information to improve target market | 3.65 | 1.17 | 3.63 | 0.81 |
| Obtain ideas from customers to improve products and services | 3.35 | 1.22 | 3.75 | 0.86 |
| People who deal with customers have information on customers and competitors | 3.94 | 0.97 | 3.63 | 0.89 |
| Alb *Strategic Development* | 3.61 | 0.57 | 3.47 | 0.99 |
| Once reviewed market research strategies are planned | 3.41 | 1.12 | 3.88 | 0.89 |
| Develop specific plans for the different market segments | 3.88 | 1.05 | 3.88 | 0.81 |
| Have the resources to improve market position | 4.06 | 1.20 | 4.19 | 0.54 |
| Competitiveness is more important than financial performance | 2.59 | 1.18 | 2.50 | 0.89 |
| Prices charged to customers are determined by customer value | 3.47 | 1.07 | 3.38 | 1.02 |
| Marketing activities are focused on markets/segments to compete effectively | 3.76 | 0.83 | 3.63 | 1.26 |
| Invest in improving market position | 4.12 | 0.99 | 4.31 | 0.60 |
| Marketing plans are according to customers and their needs rather than products/services held | 3.59 | 0.80 | 3.75 | 0.86 |
| Alc *Use of customer information* | 3.80 | 0.54 | 3.58 | 1.00 |
| Keep promises made to customers | 4.24 | 0.75 | 4.13 | 0.81 |
| Respond to customer needs when contracts are written and terms of sale are created | 3.53 | 1.07 | 3.50 | 0.73 |
| Payment terms or credit policies with clients take their needs into account | 3.71 | 0.92 | 3.25 | 1.13 |
| Respond to customer needs by delivering on time | 4.00 | 0.94 | 4.06 | 0.85 |
| **A2 Corporate Entrepreneurship** | 3.17 | 0.99 | 3.38 | 1.13 |
| Willing to take more risks than competitors | 2.82 | 1.59 | 3.31 | 1.30 |
| Better innovators than competitors | 2.94 | 1.43 | 3.38 | 1.15 |
| Plan strategies better than competitors | 3.06 | 1.39 | 3.63 | 0.72 |
| Compared to competitors, are able to identify customer's needs and wants more accurately | 3.18 | 1.33 | 3.50 | 0.89 |
| Able to make the vision of the business a reality | 3.76 | 0.66 | 4.00 | 1.21 |
| Able to identify new market opportunities better than competitors | 3.24 | 1.15 | 3.75 | 0.93 |
| **A3 Innovativeness** | 2.96 | 0.88 | 3.13 | 1.29 |

|  |  |  |  |  |  |
| --- | --- | --- | --- | --- | --- |
|  | First to market with innovative new products and services | 3.18 | 1.13 | 3.31 | 1.14 |
|  | First to develop processes that are technologically more advanced than competitors | 2.82 | 1.01 | 3.25 | 1.29 |
|  | First to recognise and develop new markets | 3.06 | 0.90 | 3.44 | 1.15 |
|  | At the leading edge of technological innovation | 2.76 | 1.09 | 3.31 | 1.01 |
| **A4** | **Learning Orientation** | 3.36 | 0.51 | 3.22 | 0.98 |
| **A4a** | **Commitment to Learning** | 3.24 | 0.58 | 3.13 | 0.98 |
|  | Management sees survival and competitiveness dependent on the ability to learn | 3.41 | 1.12 | *3.88 | 1.09 |
|  | Basic values emphasise the role of learning as a key to improvement | 3.53 | 1.07 | 3.63 | 1.09 |
|  | Employee learning is an investment, not an expense | 3.76 | 1.15 | 3.69 | 1.35 |
|  | Employee learning isn't really a top priority (reversed scoring) | 2.24 | 1.15 | 2.13 | 1.15 |
| **A4b** | **Open-Mindedness** | 3.25 | 0.54 | 3.22 | 0.95 |
|  | Not afraid to question the shared assumptions made about approach to business | 3.35 | 0.79 | *3.75 | 0.77 |
|  | Managers do not want their "view of the world" to be questioned (reversed scoring) | 2.29 | 0.69 | 2.38 | 0.89 |
|  | Employees are encouraged to be open-minded and to "think outside the box" | 3.94 | 1.03 | 3.69 | 1.14 |
|  | An emphasis on constant innovation is not part of our corporate culture (reversed scoring) | 2.41 | 1.37 | 2.44 | 1.21 |
| **A4c** | **Shared Vision** | 3.59 | 0.58 | 3.29 | 1.16 |
|  | Top leadership share its vision for the business unit with everyone in the company | 4.18 | 0.95 | 3.94 | 1.12 |
|  | Employees understand where the business unit is going | 3.76 | 0.75 | 3.56 | 0.81 |
|  | All employees are committed to the goals of this business unit | 3.29 | 1.10 | 3.25 | 1.13 |
|  | Employees view themselves as partners in charting the directions of the business unit | 3.12 | 0.93 | 3.25 | 1.18 |
| **A5** | **Technology Orientation** | 3.31 | 1.05 | 3.49 | 1.40 |
|  | New products are always state-of-the-art technology | 3.47 | 1.18 | 3.63 | 1.20 |
|  | Technological innovation based on research results are easily accepted | 3.24 | 1.30 | 3.75 | 1.13 |
|  | Technological innovation is easily accepted in programmes/project management | 3.29 | 1.21 | 3.88 | 1.36 |
|  | Sophisticated technologies are used in new product development | 3.24 | 1.20 | 3.56 | 1.31 |
| **B** | **Business Performance in Relation to its competitors** | 3.20 | 1.41 | 3.12 | 1.26 |
|  | Company's year on year on sales turnover | 3.06 | 1.85 | 3.38 | 1.15 |
|  | Division's performance year on year based market share | 3.24 | 1.25 | 3.31 | 0.87 |
|  | Division's performance year on year is based on profitability | 3.29 | 1.53 | 3.25 | 1.24 |
| **C** | **Top Management Emphasis** | 3.91 | 0.48 | 3.56 | 1.14 |
|  | We must adapt to market trends for the survival of the business unit | 3.71 | 0.77 | 3.50 | 1.26 |
|  | We must be sensitive to activities of our competitors. | 3.35 | 1.11 | 3.56 | 1.36 |
|  | We must gear up to meet customers' future needs. | 4.18 | 0.64 | # 3.88 | 0.72 |
|  | Serving customers is the most important aspect of our business unit | 4.41 | 0.62 | 4.19 | 0.83 |

\*Increased with strategic leadership coaching; #decreased with strategic leadership coaching; n=17, ± standard deviation, p-value < 0.05

The intervention group returned for a third visit, the following analysis investigates the intervention group across these three time points.

#### Iowa gambling task

Behaviourally, IGT performance showed improvements in long term consequence of outcomes for 1^st^ and 2^nd^ of the 100 trial intervals (1^st^ 100 trials Chi Sqr_(15,2)_=6.93, p=0.031; 2^nd^ 100 trials Chi Sqr_(15,2)_=6.81, p=0.033) and for all 300 trails (All 300 trials, Chi Sqr_(15,2)_=12.55, p=0.0018). The improvement was seen from time point 1 to time point 2 (1^st^ 100 trials z_n16_=2.53, p=0.011; 2^nd^ 100 trails z_n16_=2.94, p=0.0032; All 300 trials z_n16_=2.99, p=0.0027) and from time point 1 to time point 3 (1^st^ 100 trials z_n15_=2.04, p=0.0240; 2^nd^ 100 trials z _n15_=2.55, p=0.010; All 300 trials z_n13_=2.76, p=0.0057). Then the IGT score for the 300 trials was found to improve over time (All 300 trials Chi Sqr_(15,2)_=6.53, p=0.038), it was not apparent between which time points, **Table 4**. Alpha frequency was found to increase from time point 1 to time point 2 for IGT over the left frontal electrode (F_3_ t_n17_=-2.34, p=0.028) and the left central electrode (C_3_ t_n17_=-2.03, p=0.044), **Table 5**.

**Table 4.** Behavioural performance of the Intervention Group over 3 time points

|  |  | Intervention Group |  |  |  |  |  |
| --- | --- | --- | --- | --- | --- | --- | --- |
|  |  | Time point 1 |  | Time point 2 |  | Time point 3 |  |
|  |  | mean | ± | mean | ± | mean | ± |
| *Iowa gambling task* |  |  |  |  |  |  |  |
| *Long-term consequence* |  |  |  |  |  |  |  |
| First 100 trials | a b c | 0.92 | 0.26 | 1.57 | 1.26 | 5.25 | 12.67 |
| Second 100 trials | a b c | 1.11 | 0.39 | 5.25 | 11.74 | 8.50 | 16.92 |
| Third 100 trials |  | 4.93 | 12.40 | 5.84 | 12.64 | 13.44 | 19.95 |
| All 300 trials | a b c | 1.14 | 0.44 | 2.49 | 3.34 | 6.55 | 13.40 |
| *Bias to infrequency loss* |  |  |  |  |  |  |  |
| First 100 trials |  | 1.61 | 0.58 | 4.87 | 11.44 | 2.42 | 2.06 |
| Second 100 trials |  | 2.02 | 1.25 | 4.60 | 7.48 | 8.32 | 25.66 |
| Third 100 trials |  | 3.18 | 4.49 | 3.12 | 5.45 | 8.64 | 25.65 |
| All 300 trials | a | 1.85 | 0.92 | 3.84 | 7.42 | 3.11 | 5.51 |
| *Score* |  |  |  |  |  |  |  |
| First 100 trials |  | -5322 | 5605 | -5897 | 6276 | -3148 | 5268 |
| Second 100 trials |  | -2436 | 4207 | -3104 | 6086 | 909 | 3672 |
| Third 100 trials |  | -5238 | 5726 | -3628 | 6113 | -1815 | 5702 |
| All 300 trials |  | -12996 | 10905 | -12630 | 13688 | -4055 | 12837 |
| *Stroop colour-word conflict task* |  |  |  |  |  |  |  |
| Response time |  | 846 | 237 | 810 | 183 | 761 | 195 |
| Omissions (%) |  | 0.29 | 0.88 | 0.10 | 0.40 | 0.10 | 0.42 |
| Inaccurated responses (%) |  | 1.57 | 2.32 | 1.67 | 2.43 | 1.88 | 2.01 |
| Grey square count |  | 19.35 | 1.22 | 19.76 | 0.66 | 19.81 | 0.75 |

*Behavioural performance of the Intervention Group (IG) over the three time points recorded. <sup>a</sup>Improvement over time, <sup>b</sup>Improvement from time point 1 to 2, <sup>c</sup>Improvement from time point 1 to 3. Data are presented as mean with ±standard deviation, $p < 0.05$ .*

**Table 5.**
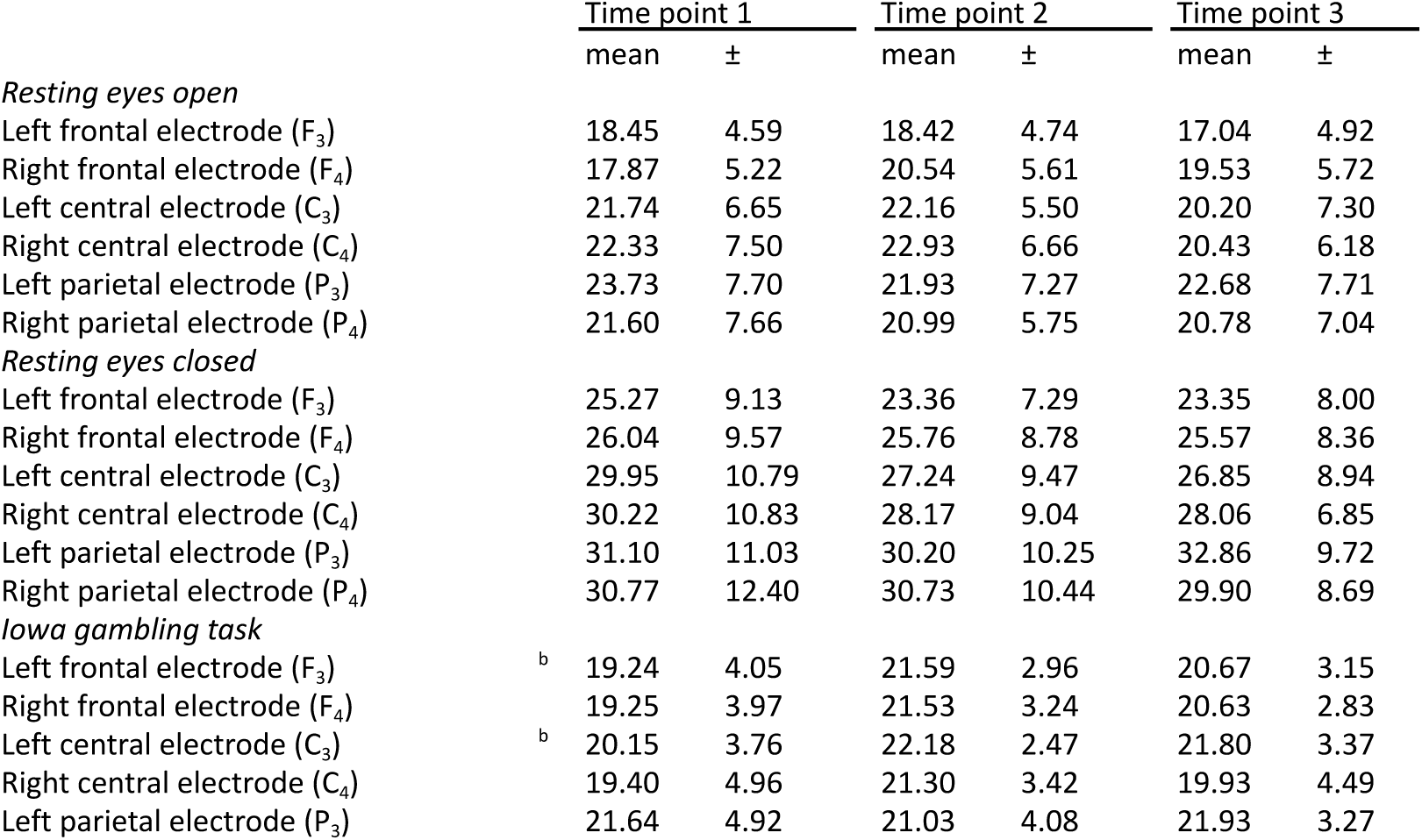
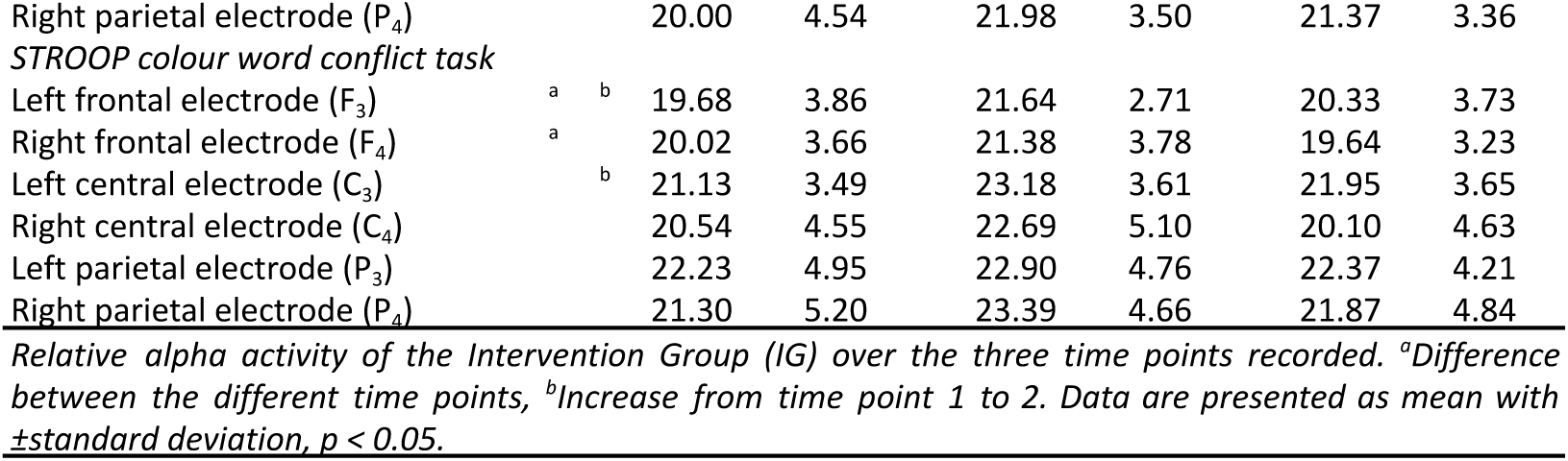
Relative alpha activity of the Intervention Group over 3 time points

#### Stroop colour word conflict task

Relative alpha frequency showed significant differences over time for Stroop task for bilateral frontal electrodes (F_3_ Chi Sqr_(17,2)_=9.88, p=0.0071; F_4_ Chi Sqr_(17,2)_=6.11, p=0.046), where left frontal relative alpha activity increased from time point 1 to time point 2 (F_3_ z_n17_=2.29, p=0.021), specific changes were not found for right frontal relative alpha activity. Then left central electrode relative alpha increased from time point 1 to time point 2 (C_3_ t_n17_=-2.05, p=0.041), **Table 5**. Event-related potential wave component differences were found for bilateral parietal electrodes, where left parietal P300 amplitude increased from time point 1 to time point 2 (P_3_ z_n17_=2.57, p=0.098) and from time point 1 to time point 3 (P_3_ z_n17_=2.15, p=0.031), and where right parietal N170 amplitude increased from time point 1 to time point 2 (P_4_ t_n17_=-2.13, p=0.034), while it decreased from time point 2 to time point 3 (P_4_ t_n17_=-2.69, p=0.015), **Figure 4**.

**Figure 4.**
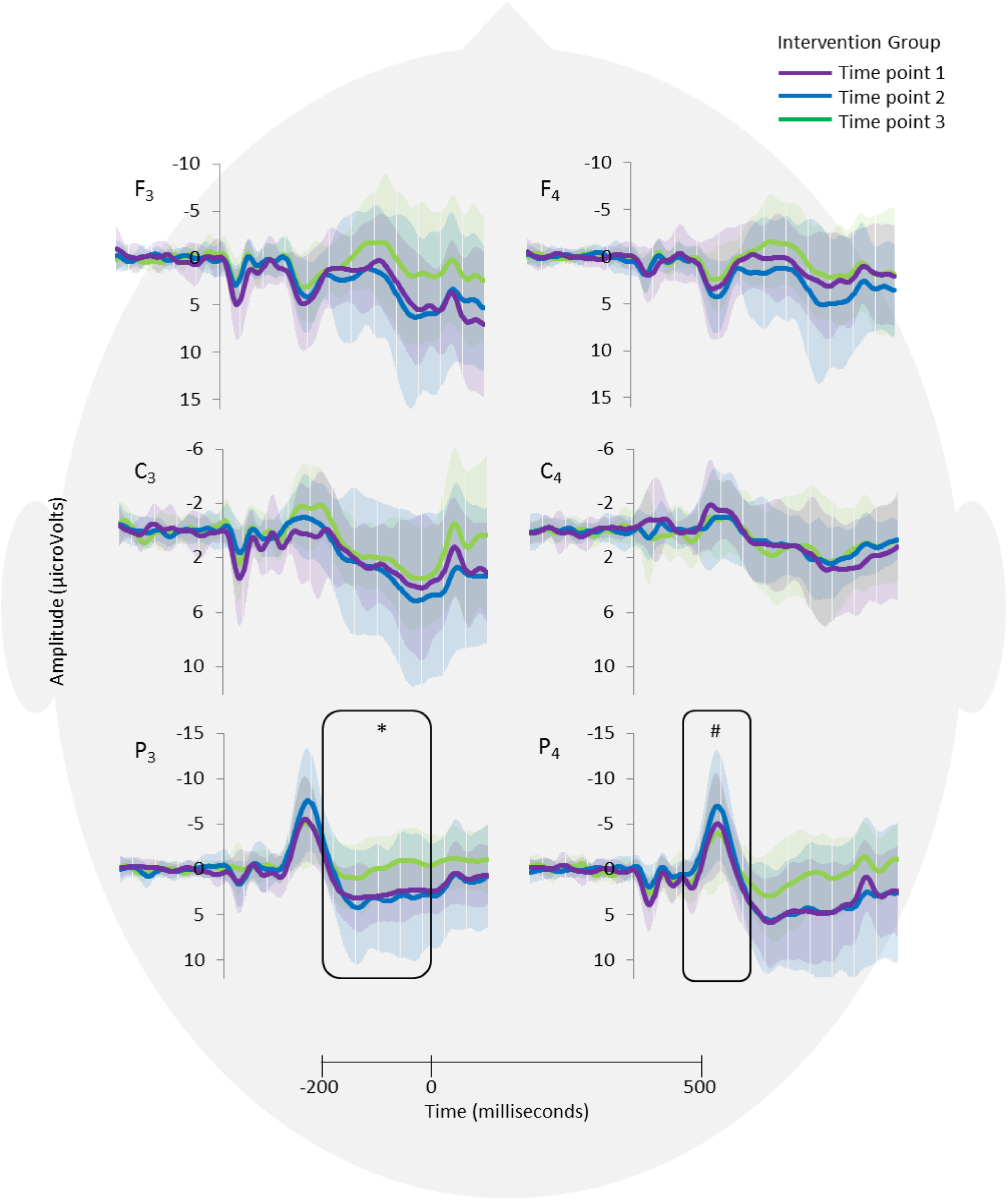
Stroop colour-word conflict event-related potentials (−200 to 500msec) for the Intervention Group (IG_n17_) at three time points, each being 2 months apart. From time point 1 to time point 2 the group completed 8 sessions of Strategic Leadership Coaching (SLC). Event-related potentials are shown for left and right frontal (F_3_, F_4_), central (C_3_, C_4_), and parietal (P_3_, P_4_) electrodes. *IG P_3_ 300 amplitude increased from time point 1 to time point 2 and 3. ^#^IG P_4_ N170 amplitude increased from time point 1 to 2 and decreased from time point 2 to 3. Data are presented as grand mean averages for each group with error bars reflecting their standard deviation.

#### STRATOR correlates

Only after coaching, time point 2, several correlates were apparent with STRATOR. During resting eyes closed left parietal relative alpha activity correlated with, subcategory of Market Orientation, Strategic Development’s “Prices charged to customers are determined by customer value” (P_3_ R_Spearman’s(n16)_=-0.80, p=0.00014). During the IGT, frontal and right central relative alpha activity correlated positively with, subcategory of Market Orientation, Strategic Implementation’s “Listen to what customers have to say” (C_4_R_Spearman’s(n16)_=0.74, p=0.00085; P_3_R_Spearman’s(n16)_=0.78, p=0.00027; P_4_R_Spearman’s(n16)_=0.75, p=0.00072), and right central relative alpha activity also positively correlated with, subcategory of Market Orientation, Strategic Implementation’s “Obtain ideas from customers to improve products and services” (C_4_R_Spearman’s(n16)_=0.74, p=0.00091). Then behaviourally, bias to infrequent loss for final 100 trials correlated negatively with subscale Top Management Emphasis’ “Serving customers is the most important aspect of our business unit” (3^rd^ 100 trials R_Spearman’s(n16)_= −0.74, p=0.00083). Then, Stroop ERP wave component, right frontal P300 latency positively correlated with subcategory of Market Orientation, Open-Mindedness (F_4_P300 latency R_Spearman’s(n16)_=-0.72, p=0.00095).

## Discussion

The main findings of this investigation support the positive effect of Strategic Leadership Coaching (SLC) on electrophysiological indices of decision-making processes, as cortical arousal improved readiness to engage with salient information which permitted stronger engagement with neural circuitry recruited for effective decision-making.

First, SLC increased alpha activity over the left hemisphere for frontal and central electrodes when young executives engaged with tasks which required activation of decision-making processes. With the 2 group comparison we saw this effect for left central electrode, **Table 2**. Within IG 3 time point analysis, this effect was repeated for left central and left frontal electrodes increase also became apparent, **Table 5**. Disappointing, was the failure of this improvement to hold two months after SLC, which suggests that young executives may require ‘top-up’ coaching sessions, **Table 5**. The total population did show improvements for frontal, central and right parietal alpha band activity however, these increases were predominantly led by IG, **Table 2**. Alpha band frequency is associated with an individual’s readiness to engage with salient stimuli (Hanslmayr *et al*., 2005; Klimesch, 1999; Klimesch *et al*., 2003; Knyazev, 2007). Our data suggests that with SLC young executives are able to recruit cognitive resource availability when they engage in cognitive tasks which require decision-making, whether assessing risk in the IGT (Bechara *et al*., 1997) or cognitive conflict in the Stroop (Stroop, 1935).

Second, SLC enhanced engagement of neural circuitry was evident, as reflected in increased amplitude of certain ERP wave components of parietal electrodes, specifically the right parietal N170 and left parietal P300 amplitudes. With the 2 group comparison, IG showed increased left parietal P300 amplitude and a tendency for increased left parietal N170 amplitude, **Figure 2**. Within IG 3 time point analysis these effects of SLC were repeated and showed significant improvement from time point 1 to time point 2, the improvement held for P300 amplitude, seen at time point 3, but did not hold for N170 amplitude, **Figure 3**. While we report improvements in amplitude for IG, CG reported decreases overtime, right central P300 amplitude decreased, **Figure 2**. The electrodes, or brain areas, involved were different for SLC increases in alpha activity and increases in ERP wave component amplitudes. Where increased alpha activity was reported frontally and centrally, while increased event-related potentials amplitude was reported parietally. Increased alpha with increased P300 amplitude has previously been reported (Intriligator and Polich, 1994; Intriligator and Polich, 1995), our findings support these and extend to include N170 wave form. Importantly, our findings suggest that the resource availability SLC created anteriorly permitted greater engagement of posterior neural circuitry involved in processing of Stroop colour-word conflict trials.

Third, to complement the electrophysiological record, behavioural performance was enhanced through SLC. During the IGT, the improvement in long term consequences was driven by the IG group, and evident 2 months post SLC. This suggests that with SLC executives learnt to play ‘lower stakes’ to ensure a more advantageous outcome overall with their decision-making, reflected mechanistically through beneficial deck choice. Behavioural response times improved, i.e. shorter response times, supporting Then during the Stroop, SLC executives improved their response times, i.e. shorter response times, supporting improved efficiency in decision making when facing cognitive conflicting information.

Prior to coaching, IG showed very strong negative relationships between the 1^st^ domain of the LSRP scale and frontal alpha activity during resting eyes closed, **Figure 3**. Limited research has been conducted to understand the relationship between alpha activity and psychopathy, the research that has been conducted reports increased psychopathy is related to decreased alpha activity (Calzada-Reyes *et al*., 2013; Ortega-Noriega *et al*., 2015; Tillem *et al*., 2019), our findings support these literature. It is unclear to why these associations were limited to IG, as there were no differences in LSRP scores between IG and CG, or why these associations were only apparent during resting eyes closed. These associations being limited to frontal cortex supports functional brain imaging studies which report dysfunctional resting state connectivity (Pujol *et al*., 2012; Yang *et al*., 2012). Importantly in the context of the current study, these associations did not confound the effects of SLC we report.

SLC allowed the young executives to set their own strategic orientation to complement their organisation. The effects of this were in part captured by administering the STRATOR, prior and post SLC, where our executives reported shifts in thinking from bottom line, i.e. financials, to the attainment of skills and new ways of thinking to promote their organisation, this included challenging accepted status quos. In addition, the executives started taking greater control over their own departments, realising they could not keep top management solely responsible for the implementation of strategy. The SLC effects were also reflected in associations with components of the STRATOR. First, positive relationships were found for relative alpha activity over central and parietal electrodes in regard to young executives taking into consideration what their customers have to say and their ideas to improve products and services, suggesting that considering customer ideas freed up cognitive resources. Second, a negative relationship was found between IGT bias infrequent loss and how they perceived top management’s view towards serving customers as the most important aspect of their unit, supporting personal responsibility over top management. Last, a positive relationship was found between right frontal P300 latency and open-mindedness to learning orientation, suggesting enhanced reflection in cognitive orientating is related to young executives’ ability to challenge status quo and seek improved ways of doing things.

Limitations to the presented study should be considered. First, this study was limited in its population selected, only recruiting male participants. Future studies will benefit from including all gender identifications, and we expect this study will support the development of these future studies. Second, our research design was lacking, and our study would have benefitted from two additions, being the inclusion of a second IG group which underwent maintenance SLC sessions (Bleich, 2016), then record of CG group for time point 3 would have supported a single model of analysis instead of the conduct of two separate analyses. With this study being unfunded, we needed to deploy a research design that was feasible and would produce results that provide a solid basic framework for future research studies.

We conclude, Strategic Leadership Coaching (SLC) improved cortical resource availability, which in turn permitted greater engagement of decision-making neural circuitry. This study provides the first and necessary neurobiological evidence to develop this line of research in SLC and in other forms of coaching, as it adds significant value.

## Acknowledgements

The authors are exceptionally grateful to the participants and their organisations for valuing this research endeavour and making this study possible.

## Author Contributions

CHN is the primary author of this study and provided coaching. KCW collected EEG measures and conducted frequency and ERP extractions. DJH set-up the testing schedule and trained KCW in methodologies. FMH designed the study, extracted behavioural performance data, applied statistical analyses, and devised the initial draft of the paper. Together all authors contributed to the manuscript substantially and supported submission for publication.

